# Gestational exposure of bisphenol-A limits decidual ECM organization via S100a10-Annexin A2 axis in murine placenta

**DOI:** 10.64898/2026.08.22.746430

**Authors:** Ankit Biswas, Somnath Mondal, Sam J. Mathew, Tushar Kanti Maiti

## Abstract

Environmental exposure to endocrine disrupting chemicals, like bisphenol-A (BPA), can impart detrimental effects on developing feto-placental unit, during pregnancy. Placenta remains a central player maintaining this feto-placental homeostasis for sustenance of a healthy pregnancy. Thus, the bisphenol-A mediated endocrine disruption affects the healthy functioning of placenta by altering key processes, such as tissue remodelling, angiogenesis, and metabolism. However, the underlying mechanism of BPA-altered ECM remodelling remains elusive. Therefore, in this study we investigated the BPA mediated changes in placental tissue remodelling using a bisphenol-A exposed murine model during pregnancy. The results reveal that, the phenotypic changes in feto-placental interface corelates with perturbed placental proteome in response to BPA. Further investigation highlights a S100a10-Annexin A2 axis mediated upregulation of tissue plasminogen activator (tPA), which drives altered extra-cellular matrix (ECM) degradation in placental decidua. This culminates into functional dysregulation in feto-placental axis, leading to reduced size of fetus and placenta. Therefore, this study provides novel insights of a S100a10-Annexin A2 axis associated mechanism for alteration of ECM remodelling in placental decidua due to BPA exposure, which may lead to toxicity related adverse pregnancy outcome.

**Graphical Abstract:** 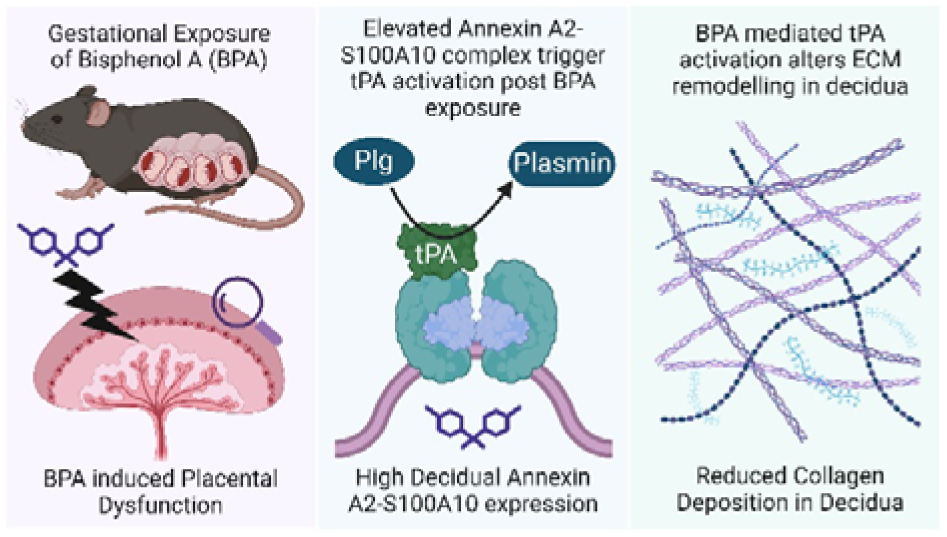

## 1. Introduction

Placenta is a dynamic organ that performs various specialized functions, which supports a healthy pregnancy through proper maintenance of the feto-maternal interface (Burton and Fowden 2015). This requires extensive structural and functional remodelling of placenta, that sustains the gestational requirements, during pregnancy (Sferruzzi-Perri et al. 2023). The placental development throughout the period of gestation (POG) is tightly coordinated by the invasion, proliferation, migration and differentiation of the trophoblast cells, which relies on the effective interaction with the extracellular matrix (ECM) constituents (Rossi et al. 2025). The complex and dynamic ECM of placenta is comprised of structural proteins, glycoproteins, proteoglycans, and matrix-associated signalling molecules, which orchestrate essential cellular function and also provides a mechanical support to the growing feto-maternal unit (Lu et al. 2026). Therefore, proper remodelling of these ECM components are essential for successful placentation.

The process of ECM remodelling is finely controlled by degradation, synthesis, and reorganization of matrix elements via the collective actions of matrix metalloproteinases (MMPs), tissue inhibitors of metalloproteinases (TIMPs), integrins, and other ECM associated proteins (Bonnans et al. 2014). These coordinated alterations drive trophoblast invasion into maternal decidua, angiogenesis, remodelling of spiral artery and feto-maternal crosstalk during early gestation. Therefore, impairment of ECM homeostasis can perturb trophoblast function during placentation, which may lead to various pregnancy complications, including preeclampsia (García-Montero et al. 2026; Famá and Pinhal 2025), fetal growth restriction (Sayres et al. 2023; Dodson et al. 2013), and preterm birth (Wang et al. 2023; Mennella et al. 2021). Increasing evidence suggests that environmental exposures may disrupt ECM organization (Wormsbaecher et al. 2020; Nikanfar et al. 2025) and cell–matrix interactions (Huang et al. 2023; Izard 2025), thereby contributing to placental dysfunction and adverse pregnancy outcomes.

Among environmental toxicants, Bisphenol-A (BPA) remains one of the abundant endocrine disrupting chemicals ubiquitously present in poly-carbonate plastics, epoxy resins, and daily consumer products. Women during pregnancy are extremely susceptible towards the toxic effects of BPA owing to its widespread environmental presence and potent activity as xeno-estrogen. Both epidemiological and experimental studies have associated prenatal BPA exposure with reproductive toxicity (Peretz et al. 2014; Cantonwine et al. 2013). Moreover, experimental murine models also demonstrated that maternal exposure to BPA is associated with detrimental effects on placenta (Adu-Gyamfi et al. 2022). This placental toxicity is mediated via altered tissue metabolism (Yue et al. 2025; Molangiri et al. 2024), aberrant angiogenesis (Tait et al. 2015; Fu et al. 2026), dysregulation of spiral artery remodelling (Müller et al. 2018), epigenetic modulation (Susiarjo et al. 2013; Ye et al. 2018) and deregulated post-translational protein modification (Biswas et al. 2026a). These functional alterations are closely associated with the maintenance of ECM homeostasis. Interestingly, Bisphenol-A is also reported to perturb ECM either via collagen interaction (Rajkumar et al. 2023) or, through regulation of ECM associated factors (Sanannam et al. 2021; Belcher et al. 2015). Although these findings demonstrate the hazardous effects of BPA on placental physiology and ECM modulation, the molecular mechanisms through which BPA alters the ECM remodelling and tissue homeostasis in placenta remain incompletely understood.

Therefore, the present study investigates the BPA mediated alteration of placental ECM homeostasis in murine pregnancy model post BPA exposure. We explored the bisphenol-A driven dynamic regulation of placental proteome to reveal higher expression of S100a10 protein, which is associated with ECM modulation. Our study demonstrated a region-specific expression of S100a10 and its interacting partners Annexin A2 and tissue plasminogen activator (tPA), which facilitates ECM remodelling required for normal placentation. BPA administration shows elevated expression of S100a10, Annexin A2 and tPA in decidua, which facilitates lower fibronectin expression and reduced collagen deposition specifically in placental decidual fraction. This finding provides a molecular mechanism for BPA associated ECM disruption in placental decidua which might contribute to altered placentation, leading to adverse pregnancy outcome.

## 2. Materials and methods

### 2.1 Animal experiments

Adult C57/BL6 mice (6 weeks, 19-25g, 10 females and 5 males) were issued from Experimental Animal Facility (EAF), Regional Centre for Biotechnology with due approval from Institutional Animal Ethics Committee (IAEC Project No. RCB/IAEC/2021/092) for conducting experiments on laboratory animals. The mice were maintained at ambient temperature with controlled environmental conditions, 12 hours of light-dark cycles, along with ad libitum access of standard chow diet and sterilized water. After a week of habituation, the females were randomly distributed in two groups. The treatment group received bisphenol-A (200 µg/kg body weight per day, dissolved in 100% ethanol) through oral gavage with corn oil for better absorption. While the mock group were gavaged with 100% ethanol in corn oil. The oral gavaging was continued for 2 weeks and then experimental mating was performed using the breeder males in 2:1 (female: male) ratio. The observance of vaginal plug next day was used for confirmation of successful mating and was inferred as e0.5 post coitus. The respective treatments were continued till e12.5, after which the pregnant dams were humanely euthanized for collection of uterine horns containing the embryos. The embryos were then dissected to collect the placental tissues for storage at -80 °C and fetal tissues for estimation of fetal parameters.

### 2.2 Differential proteomics

The placental tissues from BPA exposed (N=8) and mock (N=8) treated group were utilized for proteomic analysis as discussed previously (Kharbanda et al. 2025; Biswas et al. 2026). Briefly, the placental tissues were cryo-grinded in liquid N_2_ and then lysed with RIPA (R0278, Sigma) buffer added with 1X halt-protease and phosphatase inhibitor cocktail (78441, Thermo Scientific). Next, the lysates were acetone (014-08681, Wako) precipitated and then redissolved in 8M urea (219-00175, Wako). Further, the protein concentration was measured using BCA protein assay kit (23225, Pierce, Thermo Scientific) and 50 µg of protein was aliquoted for reduction (10 mM DL-Dithiothreitol [D5545, Sigma] for 1 h at 56 °C) and alkylation (20 mM Iodoacetamide [I1149, Sigma] in dark at RT for 1 hour). Then, the proteins were used for overnight trypsin digested utilizing sequencing grade trypsin (1:20 w/w in 50 mM ABC) (90057, Pierce, Thermo Scientific) at 37 °C. The tryptic peptides were finally desalted with Pierce C18 pipette tips (87784, Thermo scientific) before mass spectrometric acquisition.

The tryptic digest was analyzed using Sciex ZenoTOF 7600 mass spectrometer coupled with waters microscale LC system (ACQUITY UPLC M-Class System) in ZenoSWATH mode. The data were acquired in triplicates by loading 500 ng digest from each replicate onto Luna 5µm C18 Micro trap column (20 mm X 0.3 mm, 100 Å, Phenomenex), with a flow rate of 5 µL/min, which was further separated over a linear gradient of increasing organic solvent using a nanoEase M/Z HSS T3 analytical column (150 mm X 300 µm, 1.8 µm, 100 Å, Waters) for 22 minutes. The acquired spectral files (.wiff) were used for de-novo library generation in Spectronaut pulsar 19 (Biognosys) against the UniProtKB mouse proteome database with isoforms (25,709 entries, February 2026). The generated library was further used for peptide and protein identification by applying default parameters. Briefly, trypsin/P was used for specific enzyme allowing 2 missed cleavages with cysteine Carbamidomethylation (+57.021464 Da) used for fixed modification and for variable modification oxidation of methionine (+15.994915 Da), and acetylation at N-terminal (+42.010565 Da) was employed. For robust identification the precursors and peptides were filtered using 1% FDR cut-off and finally the extracted ion chromatogram (XIC) intensity of peptides were summarized for estimation of normalized protein abundance, which was further utilized for downstream analysis in R (v4.3.1) to calculate the fold change (FC) and p-values. The raw p-values were FDR adjusted using Benjamini-Hochberg (BH) correction for q-value calculation and statistical significance evaluation with a q-value cutoff of ≤ 0.05 (5% false discovery rate). Whereas the q-value filtered proteins were further evaluated for differential regulation using absolute fold change cut-off of ≥ 1.3-fold.

### 2.3 Cell culture and treatment

The HTR8/SVneo cells, which was an immortalized first-trimester extravillous trophoblast (EVTs) cell, were kept in Roswell Park Memorial Institute (RPMI) 1640 medium (R6504, Sigma) added with 2g/L sodium bicarbonate (40151UR-K05, SDFCL), 100 units/mL penicillin, 100 g/mL streptomycin (A001A, Himedia) and 10% heat-inactivated fetal bovine serum (FBS) (A5256501, Gibco). The cells were maintained at 37 °C, with 5% CO_2_ in a CO_2_ incubator under humidified conditions. Cells were passaged at least three times post revival, followed by seeding into a 6 well plate (3X10^5^ cells/well). After reaching an optimal confluency the cells were subsequently serum-starved with 1% FBS-supplemented incomplete medium for 4 hours and then exposed to 10 µM bisphenol-A (239658, Sigma) dissolved in DMSO (472301, Sigma) for 24 hours. Cells subjected to 24 hours of 0.01% DMSO treatment was designated as mock control.

### 2.4 Immuno-blot assay

The harvested placental tissue and EVT samples post BPA treatment were homogenized in RIPA (R0278, Sigma) lysis buffer for lysate preparation. Next, 30 µg of proteins from each lysate were loaded onto 12% tricine-SDS-polyacrylamide gel for electrophoretic separation at 100 volts. Subsequently, the resolved proteins were transferred to 0.22 µ polyvinylidene fluoride (PVDF) membrane (GVWP14250, Millipore, Merk) and then blocked with 5% (W/V) skimmed milk (GRM1254, Himedia) for 1 hour. Finally, the respective blots were incubated with primary antibodies against S100a10 (1:1000, 11250-1-AP, Proteintech); Annexin A2 (1:5000, 60051-1-Ig, Proteintech) and GAPDH (1:10000, A19056, ABclonal) at 4 °C for overnight. Next day, the membranes were washed with tris-buffered saline-tween (TBST) (0.1% [vol/vol]), followed by incubation in HRP conjugated secondary anti-rabbit or anti-mouse antibody (1:10000 & 1:10000 respectively) at room temperature. After an hour, the blots were again washed in TBST and then developed using Immobilon Forte Western HRP Substrate (WBLUF, Millipore, Merk) in Image Quant LAS 4000 (GE Healthcare Bio-Sciences AB, Sweden) instrument. The respective band intensities were estimated using Fiji (v2.17.0) software, which was then normalized with GAPDH abundance to determine the relative protein expression.

### 2.5 Immuno-fluorescent analysis of cells and tissue sections

The EVTs, seeded on coverslips, were treated with BPA and then fixed in 4% paraformaldehyde, post-treatment. Next, the coverslips were immuno-stained with primary antibodies against S100a10 (1:250, 11250-1-AP, Proteintech) and Annexin A2 (1:750, 60051-1-Ig, Proteintech) at RT for 1.5 hour, followed by washing and secondary antibody incubation (1:500, Alexa fluor-594 goat anti-mouse lgG and Alexa fluor-684 goat anti-rabbit lgG, Thermo Fisher Scientific) at RT for an hour. Finally, DAPI staining was performed and then the slides were imaged using a confocal microscope (Leica TCS SP8).

Harvested placental tissue was embedded in Cryomatrix (P0091, Sigma) cryo-sectioning medium. The embedded tissue was sectioned at 10 um thickness using a cryo-microtome (Slee), and adjacent sections were collected on charged glass slides (631-0108, Avantor). The sectioned slides were then washed with PBS, and the antigen retrieved in citrate buffer (1.8 mM citric acid and 8.2 mM sodium citrate in water). The sections were blocked using TNB blocking buffer (FP1012, Perkin Elmer) for 1 hour at RT. This was followed by incubation overnight with primary antibodies against S100a10 (1:500, 11250-1-AP, Proteintech), Annexin A2 (1:750, 60051-1-Ig, Proteintech), tPA (1:500, 10147-1-AP, Proteintech) and Fn1 (1:400, 15613-1-AP, Proteintech) diluted in TNB buffer at 4 °C. The signal was enhanced for S100a10-Annexin A2 and tPA-Annexin A2 co-staining using biotin-streptavidin incubation overnight, for the respective different sets of slides, and signal amplification was performed using Vectastain (PK□ 6100, Vector Laboratories) and TSA reagent kits (NEL741B001KT, NEL744B001KT, Perkin Elmer). The stained sections were imaged using a Nikon Eclipse Ti fluorescence microscope with a DS□ Qi2 camera. Each image was maximally projected; fluorescence intensity and Pearson correlation coefficient were measured using Fiji (v2.17.0).

### 2.6 Sirius Red staining

To study collagen deposition, the placental sections were stained with Sirius red. Briefly, sections were fixed for 1□h in Bouin’s fixative (25% Formaldehyde, 75% saturated picric acid, and 5% Glacial acetic acid) at 56°C, followed by washing with tap water, and stained with Picro□ Sirius red (365548, Sigma) for 1.5□h, and then washed with 0.5% acetic acid. The sections were dehydrated in 50%, 70%, and 100% ethanol, equilibrated in xylene, and mounted in DPX (06522, Sigma). The sections were imaged using a Nikon Eclipse Ti microscope with a DS□ Fi2 camera and quantified by using Fiji (v2.17.0).

### 2.7 Bioinformatic and statistical analysis

Data were plotted and graphs were generated using Prism 8 software (GraphPad Software, USA) or ggplot (3.5.1) package in R (v4.3.1), if not either mentioned. The upset plots and heat-maps were created utilizing UpsetR (1.4.0) and pheatmap (1.0.12) package respectively. Pathway enrichment was performed via clusterProfiler (v4.10.1) package using ReactomePA (v1.46.0) database. All the graphical illustrations were made in BioRender (https://www.biorender.com/) and the image analysis was performed using Fiji (v2.17.0).

The data were plotted for at least three biological replicates and expressed as mean ± Standard error of mean (SEM) in Prism 8 software (GraphPad Software, USA), followed by statistical testing post-treatment using Student’s t-test, where the P values ≤ 0.05 were considered as statistically significant and are indicated by asterisks as follows: *P ≤ 0.05, **P ≤ 0.01, ***P ≤ 0.001, ****P ≤ 0.0001.

## 3. Results

### 3.1 Bisphenol-A drives feto-placental alteration through placental proteome perturbation

Bisphenol-A is a xeno-estrogen, which is abundantly present in routinely used household items. Its toxic exposure during pregnancy can exert detrimental effect on the developing feto-placental interface. Therefore, to decipher the BPA mediated toxicity mechanism, we developed a murine model through gestational exposure to bisphenol-A, according to previously established protocols (Mao et al. 2020; Biswas et al. 2026a). **Figure 1A** depicts the schematic pipeline followed for model generation and subsequent assessment of feto-placental parameters. The assessment revealed no significant changes in the number of embryos, weight of fetus and placental weight (**Figure 1B, C and I**). However, the BPA exposure led to significant reduction in CRL (crown-to-rump length) (**Figure 1D-E**), APD (abdominal antero-posterior diameter) (**Figure 1F-G**) and placental diameter (**Figure 1J-K**). Interestingly, feto-placental weight ratio exhibited significant elevation but with limited effect size (**Figure 1H**). Therefore, to determine the underlying mechanism of these gross alterations, we performed proteomic analysis of placental tissues from BPA exposed and mock treated group.

**Figure 1:**
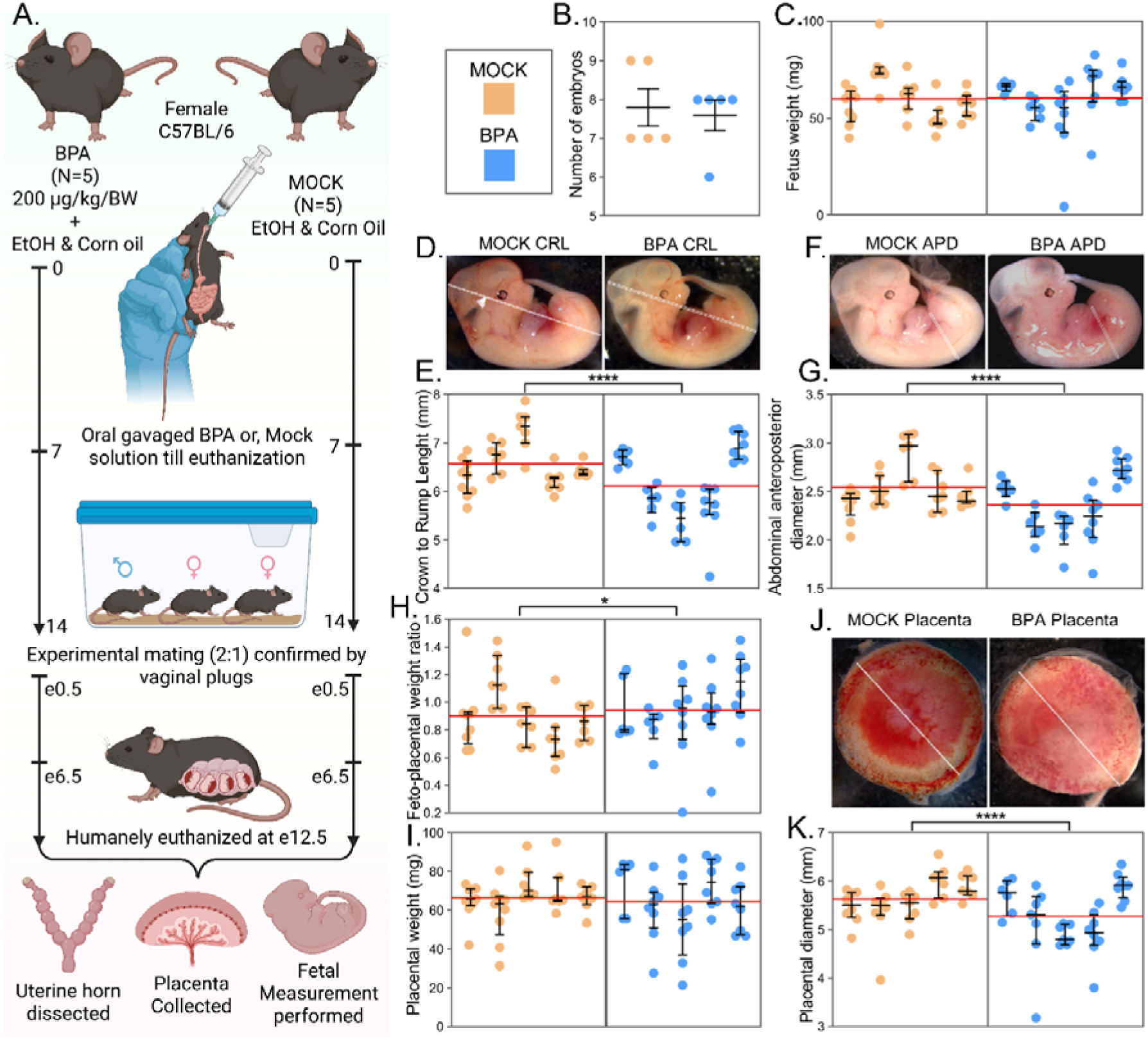
Phenotypic assessments of feto-placental parameters post BPA administration. **A)** The schematic pipeline followed for development of bisphenol-A exposed murine model during pregnancy. The measurement of **B)** embryo number, **C)** fetal weight, **E)** crown-to-rump length (CRL), **G)** abdominal antero-posterior diameter (APD), **H)** feto-placental weight ratio, **I)** placental weight and **K)** placental diameter at embryonic day 12.5 from both the mock treated and BPA exposed group. The representative image of fetal **D)** CRL, **F)** APD and placental **J)** diameter marked in white from both the comparison group. Individual litter data are shown as mean ± SEM (black lines and error bars), while the red line denotes the overall mean across all litters **(n=5)**. Differences between groups were evaluated using nested t-test., considering p-value≤0.05 as significant.

We followed the experimental design depicted in **figure 2A**, to map the murine placental proteome landscape, which identified 35,250 peptides corresponding to 3982 protein groups. Further data exploration showed, no clear separation between BPA and mock group in principle component analysis (PCA) (Figure 2B), but the median coefficient of variation (CV) of BPA (24.4%) and mock (28.1%) group reflected a considerable biological variation sufficient for differential analysis (**Figure 2C**). Next, we performed differential protein expression analysis between BPA and mock group to identify 35 upregulated and 31 downregulated proteins using absolute log2 FC cutoff of ≥ 0.378 and FDR adjusted p-value (q-value) cutoff of ≤ 0.05 (Figure 2D). The heat-map highlight the regulated proteins which clusters perfectly according to their regulatory direction (**Figure 2E**). The pathway enrichment by gene set enrichment analysis (GSEA) using Reactome database displayed the upregulation of pathways associated with metabolism, apoptosis and signal transduction. Meanwhile, the pathways related to mitotic pro-phase, translation, vesicle mediated transport, metabolism of proteins and post-transnational protein modification were downregulated (**Figure 2F**).

**Figure 2:**
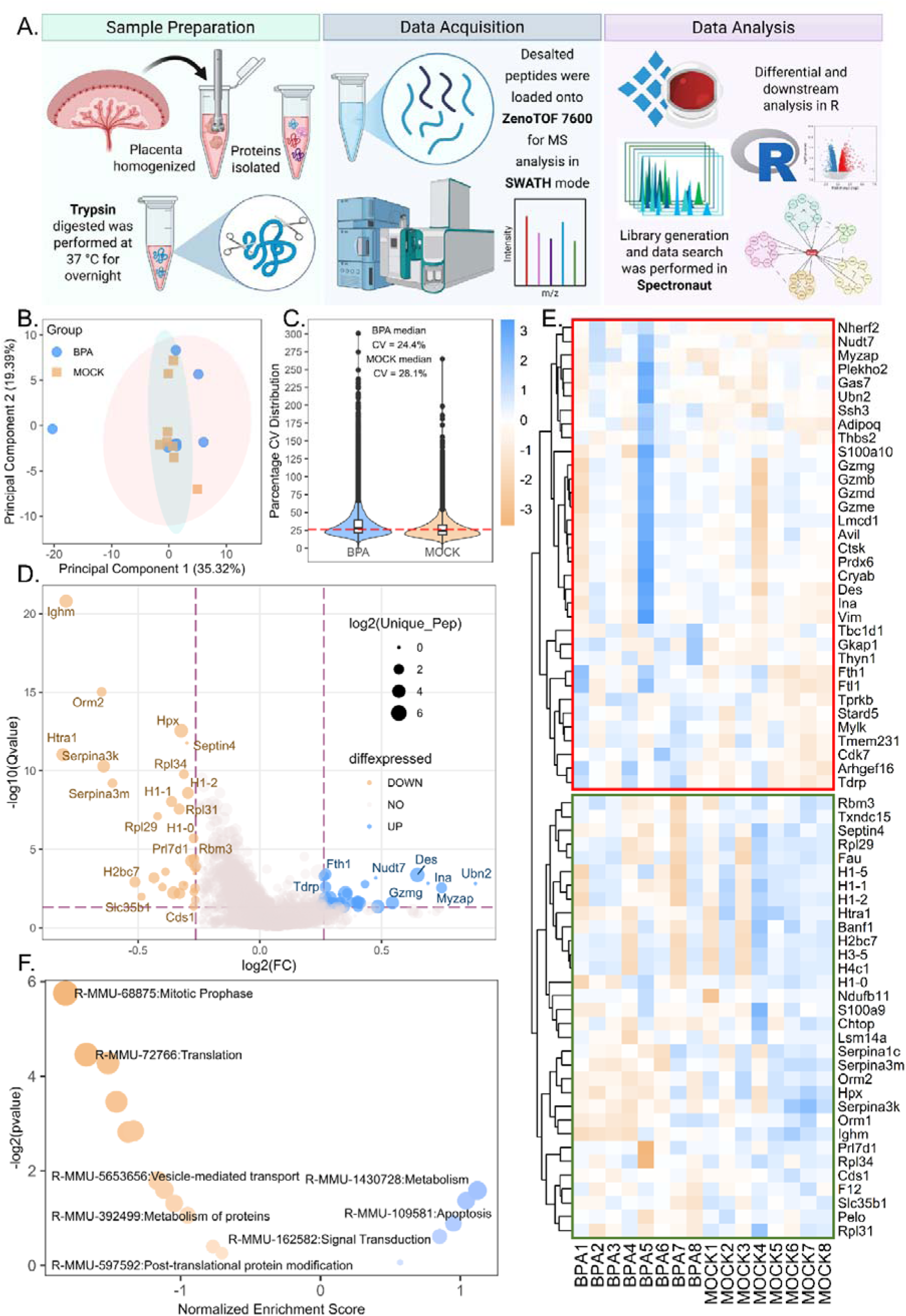
Bisphenol-A drives placental proteome signature modulation. **A)** The workflow adopted for proteome profiling of murine placenta from mock treated and BPA exposed dams. **B)** The principal component analysis grouped the samples from each comparison set using PC1 in x-axis and PC2 in y-axis. **C)** The violin plot displays percentage coefficient of variation (CV) distribution for BPA and mock group. **D)** The volcano plot shows the differentially expressed proteins in BPA vs mock comparison **(n=8)**, where x-axis represents log2 fold change (FC) and y-axis represents -log10 FDR adjusted p-value (Qvalue). The dotted lines denote the differential cut-off, while the circle size refers to number of unique peptides for each identified proteins. **E)** The heatmap indicates the regulation pattern of 67 differentially expressed proteins (DEPs), where the red square denotes upregulated cluster and the green square refers to downregulated cluster. **F)** The volcano plot shows the DEPs enriched Reactome pathways with normalized enrichment scores in x-axis and -log2 p-value in y-axis.

### 3.2 BPA elicits comparable dysregulation on both trophoblast and placenta proteome

The tissue architecture of placenta was heterogeneous; therefore, the effects of bisphenol-A would widely vary depending on cell type. Thus, we utilized our previously published proteome data (Biswas et al. 2026a) of BPA exposed extravillous trophoblast cells (EVTs) to characterize comparable changes across cellular and tissue levels. The placenta data enriched 40 pathways, whereas the EVTs data showed enrichment of 103 pathways. The comparison between these regulated pathways indicated a considerable overlap of 21 Reactome enriched pathway, that displayed both concordant and opposing patterns of regulation (**Figure 3A**). Despite this observation, the cnet plot revealed an association of completely different set of proteins with the overlapped pathways from both the EVTs (**Figure 3C**) and placenta (**Figure 3B**) data, which led us to compare the differentially expressed proteins (DEPs) from EVTs and placenta groups. The placenta and EVTs data identified 67 and 83 DEPs respectively, but surprisingly only 2 DEPs showed overlap (**Figure 3D**). Among them, Histone H4 (H4c1) showed upregulation in EVT samples but revealed decreased expression in placental data. However, the other protein, S100a10 displayed significant upregulation in both EVTs and placental samples post BPA exposure (**Figure 3E**).

**Figure 3:**
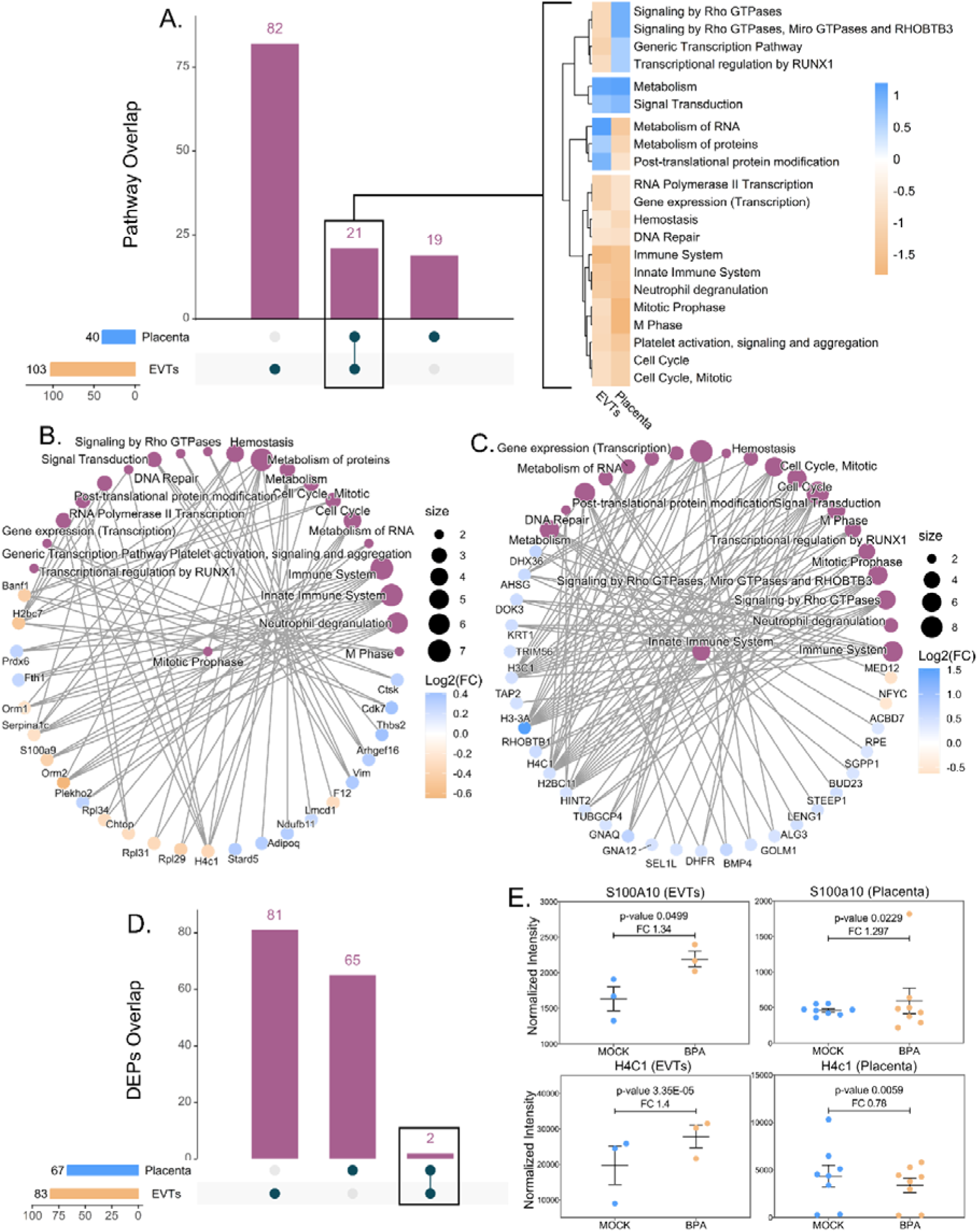
Functional overlap between BPA-altered EVTs and placental proteome. **A)** The upset plot highlights the overlap between pathways enriched from BPA exposed EVTs and placenta proteome data. The common pathways are plotted in a heatmap showing the regulatory direction across EVTs and placenta samples. The category network (cnet) plot of overlapped pathways displays pathway-protein association from **B)** placenta and **C)** EVTs data. In the network, protein nodes are colour-coded by log2 fold change (FC), and the pathway node size corresponds to the number of associated proteins. **D)** Upset plot shows the intersection between differentially expressed proteins (DEPs) from EVTs and placenta proteome dataset. **E)** The normalized intensity of the common proteins is plotted in a box-plot for EVTs and placenta data separately.

### 3.3 Bisphenol-A administration exhibits elevated expression of S100a10 and Annexin A2 in placenta

The EVTs and placenta proteome data revealed the BPA mediated upregulation of S100a10 levels, which functions in complex with Annexin A2. This S100a10 and Annexin A2 complex perform ECM remodelling through fibrinolysis, plays role in membrane trafficking and modulates ion channel localization (Okura et al. 2023). Therefore, we investigated the regulation and localization of both S100a10 and Annexin A2 proteins in EVTs and placenta samples post BPA exposure (**Figure 4A**). The western blot analysis validated the bisphenol-A driven upregulation of S100a10 and Annexin A2 in both EVTs and placenta lysate (**Figure 4B**). The densitometric analysis of the blots showed statistical significance between the mock and BPA group (**Figure 4C**). However, immunofluorescent staining of EVTs post BPA exposure did not exhibit any defect in localization of S100a10 and Annexin A2 proteins. Moreover, the BPA treatment also did not disrupt the co-localization of S100a10-Annexin A2 complex (**Figure 4D**).

**Figure 4:**
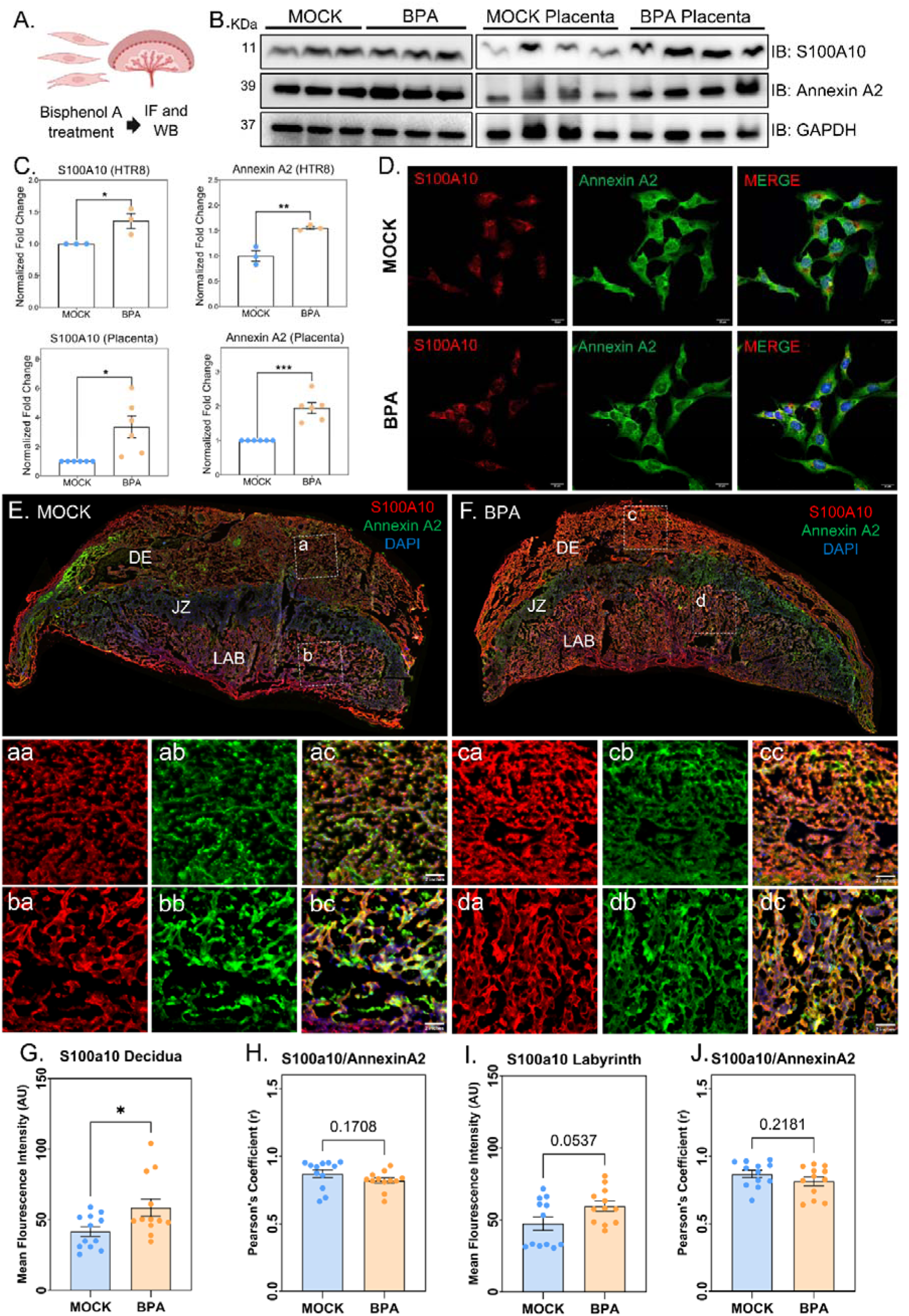
Bisphenol-A triggered dysfunction across S100a10-Annexin A2 axis. **A)** general experimental scheme utilized for validation study. **B)** Western blot detection of S100a10, Annexin A2, and GAPDH in the mock-treated and bisphenol-A exposed lysates from both EVTs **(n=3)** and placenta **(n=6)** group. **C)** The densitometric estimation of GAPDH normalized fold change for S100a10 and Annexin A2 is summarized in a box-plot. **D)** The BPA exposed and mock treated EVTs are immune-stained for S100a10 (red), Annexin A2 (green) and DAPI (blue). Scale bar = 20 µm. The representative immunofluorescence micrographs of transversely sectioned placental tissues from **E)** mock treated and **F)** BPA-exposed samples **(n=4)**, which are immune-stained for S100a10 (red), Annexin A2 (green) and DAPI (blue). The inset displays corresponding images for mock decidua (**aa, ab, ac**), & labyrinth (**ba, bb, bc**), and BPA-exposed decidua (**ca, cb, cc**), & labyrinth (**da, db, dc**). Estimation of mean fluorescence intensity (AU) of S100a10 from mock and BPA-treated **G)** decidua, & **I)** labyrinth, and the Pearson correlation coefficients for S100a10-Annexin A2 colocalization in **H)** decidua and **J)** labyrinth. Data is plotted as mean ± SEM. Student’s t-test was performed, with p≤0.05 considered significant. Scale bar = 2 inches (ac, bc, cc, dc).

Next, we immuno-stained the placental sections, which interestingly showed region specific staining for S100a10, but not for Annexin A2 in both mock treated and BPA exposed placental sections (**Figure 4E-F**). The S100a10 staining revealed significantly higher expression in placental decidua (**Figure 4aa, ca and G**) and labyrinth (**Figure 4ba, da and I**) post BPA treatment, whereas no expression was observed in junctional zone. Moreover, Annexin A2 demonstrates no significant modulation of distribution across different layers of placenta after BPA exposure (**Figure 4ab, bb, cb and db**). Further analysis confirmed that, BPA treatment did not affect the co-localization of S100a10 and Annexin A2 in both decidua (**Figure 4ac, bc and H**) and labyrinth (**Figure 4cc, dc and J**) layer of placenta, as observed in case of EVTs also.

### 3.4 Higher expression of decidual tPA impairs fibronectin and collagen deposition in placental decidua due to BPA exposure

The S100a10 and Annexin A2 complex recruits both tissue plasminogen activator (tPA) and plasminogen to facilitate the conversion of plasmin, which activate the matrix metalloproteinases for fibrinolysis and ECM remodelling (Hajjar et al. 1998; Surette et al. 2011). Therefore, we next elucidated the expression of tPA in different layers of placental sections and checked its spatial co-localization with Annexin A2 using immuno-staining. The analysis revealed selective abundance of tPA across different regions of placenta from mock treated and BPA exposed groups (**Figure 5A-B**). Further exploration highlighted increased tPA expression in both decidual (**Figure 5aa, ca and C**) and labyrinth (**Figure 5ba, da and E**) fraction of the placenta due to BPA exposure. However, the Annexin A2 expression remained unchanged as observed previously (**Figure 5ab, cb, bb and db**). Interestingly, BPA treatment did not alter the decidual co-localization of tPA and Annexin A2 (**Figure 5ac, cc and D**) but significantly reduced the spatial overlap in labyrinth region (**Figure 5bc, dc and F**).

**Figure 5:**
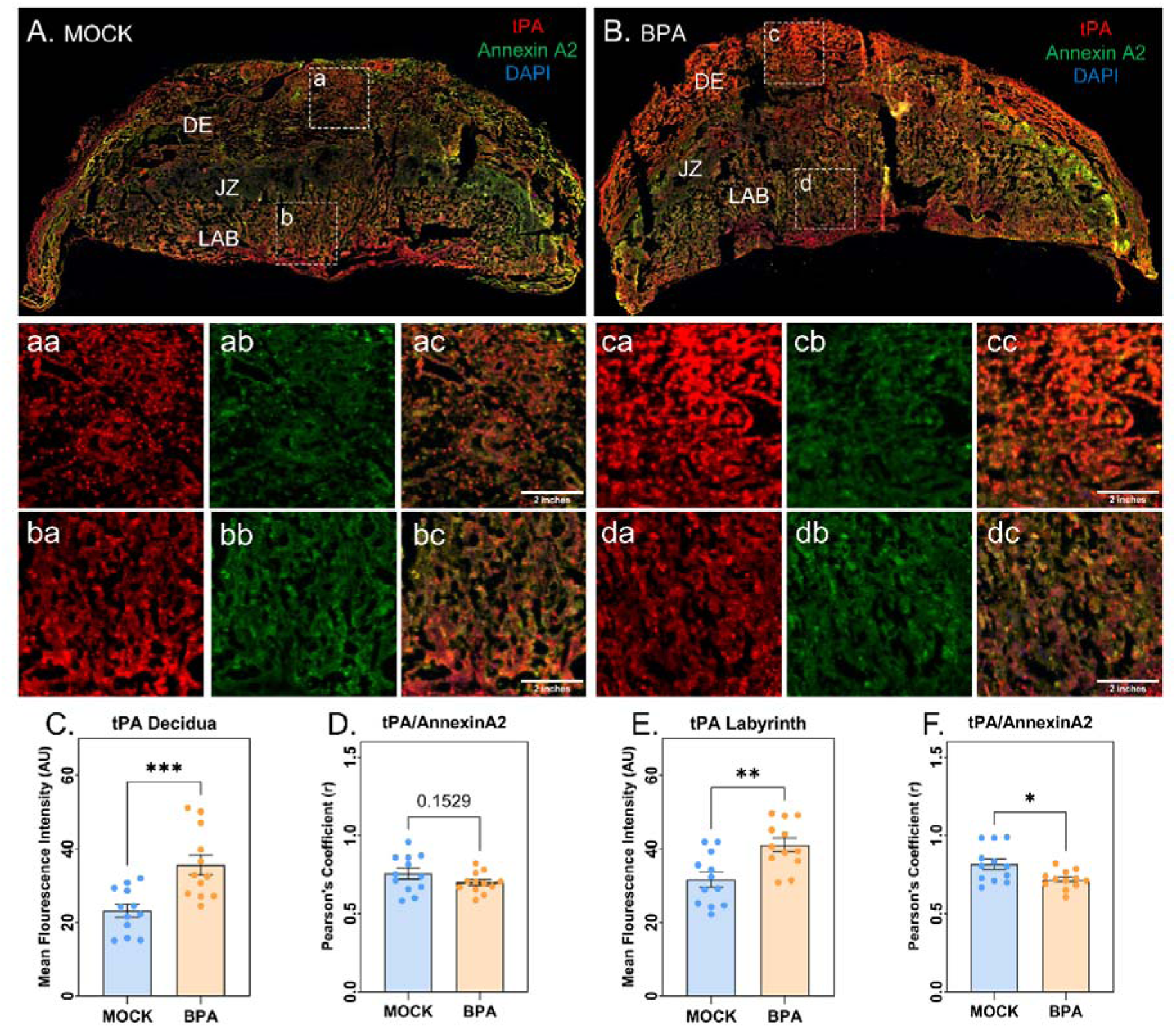
BPA exposure results in increased placental tPA levels. Representative immunofluorescence micrographs of transverse placental sections from **A)** mock and **B)** BPA exposed dams, stained for tPA (red), Annexin A2 (green), and DAPI (blue). The insets show representative images of the mock decidua (**aa, ab, ac**), & labyrinth (**ba, bb, bc**), and BPA-treated decidua (**ca, cb, cc**), & labyrinth (**da, db, dc**). Quantification of mean fluorescence intensity (AU) of tPA from mock and BPA-treated **C)** decidua and **E)** labyrinth, and the Pearson correlation coefficients for tPA and Annexin A2 colocalization in **D)** decidua and **F)** labyrinth. Data is represented as mean ± SEM **(n=4)**. Student’s t-test was performed, with p≤0.05 considered significant. Scale bar = 2 inches (ac, bc, cc, dc).

These results suggested that BPA triggers elevated expression of S100a10 which recruit more tPA in decidual layer of placenta. These higher tPA expression would ultimately culminate in impaired ECM remodelling. Therefore, we next explored the abundance of Fibronectin (Fn1), a major ECM protein and also evaluated the status of collagen deposition in different placental layers, post BPA administration. The immuno-staining of Fn1 indicated its distinct spatial expression in mock and BPA exposed placental sections (**Figure 6A-B**). Further assessment revealed a lower fibronectin expression in decidua (**Figure 6a, c and C**) but its levels were elevated in placental labyrinth regions (**Figure 6b, d and D**) due to BPA exposure. Next, we stained the placental sections with Sirius red to assess the collagen deposition. The Sirius Red staining displayed collagen abundance in red across the layers of decidua and junctional zone, however the labyrinth region shows no positive staining for Sirius red in both mock and BPA exposed samples (**Figure 6E-F**). Further estimation showed, a significant reduction in collagen positive area across placental decidua post BPA treatment, although the collagen abundance in junctional zone remained unchanged (**Figure 6G-I**).

**Figure 6:**
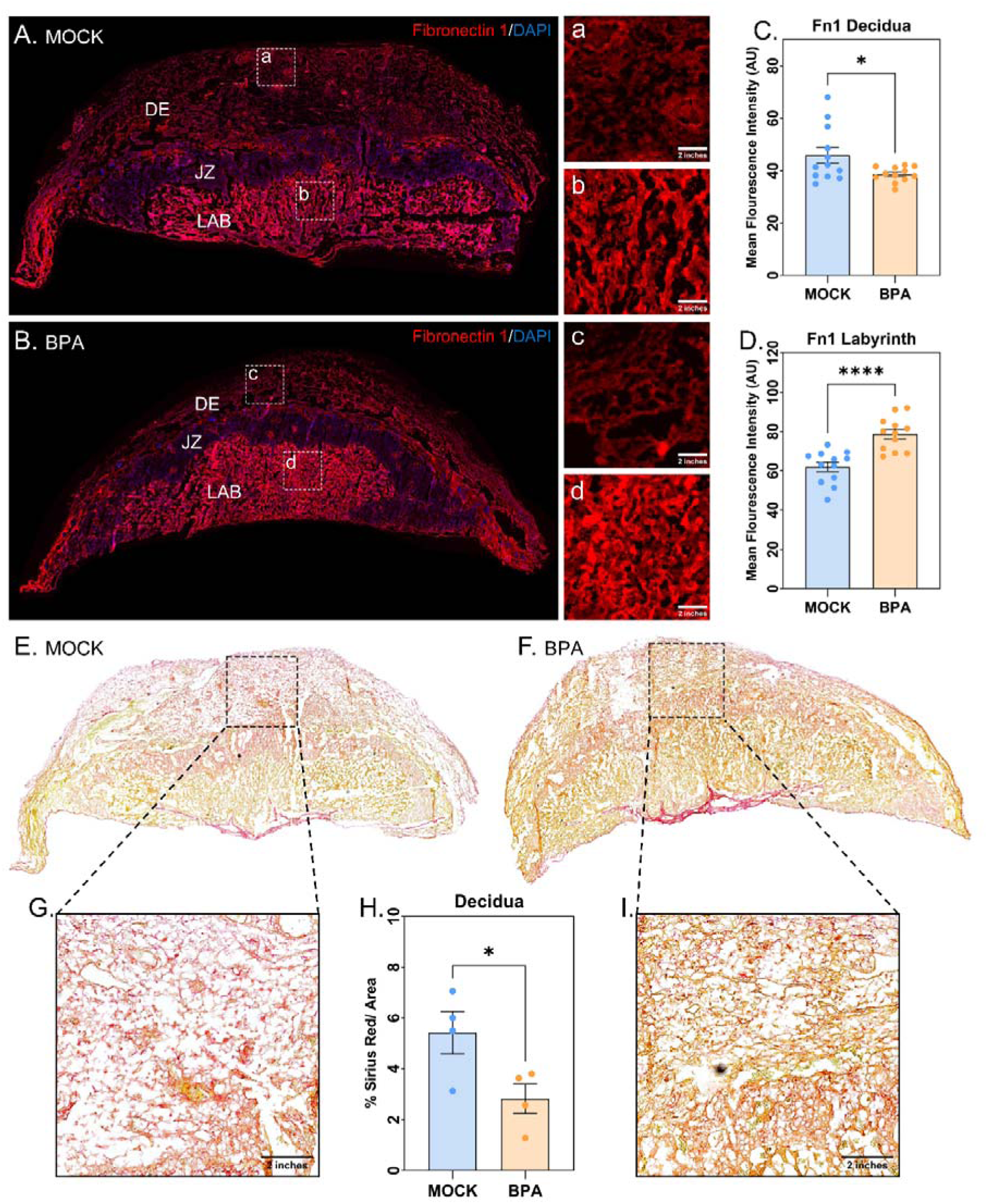
Bisphenol-A remodels decidual extracellular matrix components in placenta. Representative immunofluorescence micrographs of a transverse section of placenta from **A)** mock and **B)** BPA-exposed dam, stained with Fibronectin 1 (red) and DAPI (blue). The inset shows representative images from the **a)** mock decidua & **b)** labyrinth, and BPA-treated **c)** decidua & **d)** labyrinth. Quantification of mean fluorescence intensity (AU) of tPA from mock and BPA-treated **C)** decidua, & **E)** labyrinth. Representative bright field micrographs of the transverse sections of placenta from **E)** mock and **F)** BPA-treated mice, stained for Sirius red to label the extracellular matrix. The inset shows of Sirius red stained decidua from **G)** mock and **I)** BPA exposed placenta. **H)** The estimation displays the percentage of Sirius red+ area as a fraction of total decidual area. Data is presented as mean ± SEM **(n=4)**. Student’s t-test was performed, with p≤0.05 considered significant. Scale bar = 2 inches (a, b, c, d).

## 4. Discussion

During gestation the pregnant women are exposed to various environmental contaminants, such as bisphenol-A which acts as potent toxicants, owing to its ubiquitous occurrence in daily consumer products. The presence of bisphenol-A has been detected in maternal serum, urine, cord blood, and placenta, in variable amounts widely depending on race, country, economical status etc. However, multiple epidemiological studies have associated higher systemic concentrations of bisphenol-A in mothers with adverse pregnancy outcomes. Such studies suggest a positive association of preterm birth with increased BPA concentration in maternal urine (Huang et al. 2019; Cantonwine et al. 2015). Similar correlation is also observed for preeclampsia where, higher amount of BPA in maternal serum (median: 3.40 vs. 1.50 µg/L, p-value ≤ 0.01) (Ye et al. 2017) and urine (Philips et al. 2019) is associated with preeclampsia outcome. Whereas, elevated bisphenol-A levels in maternal urine, plasma and cord blood are correlated with lower birth weights in neonates (Chou et al. 2011; Li et al. 2024).

Therefore, several studies have developed BPA exposed murine models to investigate the toxic effects of BPA, that leads to adverse pregnancy outcome (Wang et al. 2025; Müller et al. 2018; Ye et al. 2018; Sun et al. 2024). However, the exposure concentration, the route of administration and the embryonic day at sacrifice differs considerably between studies. Thus, we have followed a previously published and validated protocol (Mao et al. 2020; Biswas et al. 2026a) to develop the exposure model in pregnant mice by administering 200 µg BPA/kg body weight orally everyday till embryonic day 12.5. Previously performed dose response studies support this bisphenol-A dosage, which is also substantially lower than the diet-administered maximum nontoxic dose for BPA in rodents (200 mg/kg body weight per day) (Dolinoy et al. 2007; Vandenberg et al. 2010). More importantly, we sacrificed the pregnant dams at mid-gestation (e12.5), as it represents the placental stage with distinct layers already formed, which is now transitioning into a stable functional phase to support the fetal growth (Bolon et al. 2025). Therefore, at this stage the endocrine disruptions, altered gene expressions and structural defects at feto-placental axis can be detected easily. Thus, here also we observed a bisphenol-A mediated gross modulations of feto-placental parameters, with significant reduction of fetal and placental size. These overall alterations indicate a potential role of bisphenol-A in placental impairment which may lead to growth defects in murine embryos. Therefore, we explored the proteome landscape of placenta to dissect the intricate BPA driven molecular changes at feto-placental interface.

The murine placenta proteome exhibits a good data quality which displays considerable modulation of protein expression across different signalling cascades. These 67 distinctly regulated proteins are involved in the alteration of specific pathways previously shown to be affected by bisphenol-A, which includes metabolism (Yue et al. 2025; Molangiri et al. 2024), signal transduction (Sun et al. 2024; Wang et al. 2025), apoptosis (Celar Sturm et al. 2025; Ponniah et al. 2015), and post-translational protein modification (Biswas et al. 2026b,a). However, the BPA triggered response in placental tissue may extensively vary depending upon the cell type heterogeneity. Therefore, we integrated our previously published BPA exposed cellular proteome data (Biswas et al. 2026a) with this tissue proteome data to capture the common regulatory signalling cascades, which may act similarly in diverse placental cell types in response to bisphenol-A. Interestingly, we observed a substantial overlap between pathways enriched from EVTs and placenta data. However, closer inspection showed that completely different sets of proteins from the EVTs and the placenta data are associated with the overlapped pathways. This suggests a presence of cell and tissue specific molecular players which are driving similar pathway level dysregulation in response to BPA exposure. Therefore, we compared the differentially expressed proteins from both the EVT and placental data to map the overlapping molecular determinants of BPA toxicity. Surprisingly, only two proteins show overlap, among which S100a10 displays significant elevation in both the placenta and EVT proteome data.

S100a10 is a small, multi-functional protein belonging to S100 family, which is involved in tissue remodelling and immune modulation (Okura et al. 2023; Saiki and Horii 2019). Despite, belonging to a calcium binding protein family, the activity of S100a10 does not rely upon calcium binding but its interaction with a cytosolic protein, Annexin A2 (Rescher and Gerke 2008). S100a10 largely remains in a heterotetrameric complex with Annexin A2, which facilitates its intracellular stability via preventing its proteasomal degradation (Bharadwaj et al. 2021). This S100a10-Annexin A2 complex interacts with various functional proteins to regulate fibrinolysis, vesicular trafficking, ion channel localization and autophagy (Okura et al. 2023; Saiki and Horii 2019). Previously, its expression was also reported in human placental syncytium, trophoblast cells, blood vessel endothelial cells and amniotic membranes (Abd El-Aleem and Dekker 2018). Therefore, we investigated the fate of S100a10-Annexin A2 complex in EVTs and placenta post bisphenol-A exposure. The immuno-blot result reveals a modest increase in S100a10 and Annexin A2 abundance in EVTs but placental tissue displays a strong upregulation for both in response to BPA. This observed increase in S100a10 and Annexin A2 expression may result from BPA mediated activation of transcription factor c-JUN (Zhang et al. 2025; Lan et al. 2017; Biswas et al. 2026a), as both of them are transcriptionally regulated via c-JUN (Moreau et al. 2015; Chottekalapanda et al. 2020). Moreover, the immuno-staining of EVTs does not shows any change in either expression or localization of the complex, which suggests that S100a10 mediated toxicity mechanism of BPA is not EVT associated. Similar to EVTs, the placental sections also did not display altered localization and change in Annexin A2 expression but, we observed a BPA triggered elevation of S100a10 protein specifically in placental labyrinth and decidua. This led us to investigate the BPA altered signalling cascades downstream to S100a10-Annexin A2 complex in placental tissues.

Bisphenol-A is known to impair placental tissue decidualization (Wang et al. 2025) and angiogenesis (Tait et al. 2015; Fu et al. 2026), which are important processes closely linked with extra-cellular matrix remodelling. Therefore, we explored S100a10-Annexin A2 mediated regulation of downstream targets, such as plasminogen and tissue plasminogen activator (tPA), which are involved in fibrinolysis and ECM remodelling. Annexin A2 anchors the heterotetrameric complex with S100a10 in cell surface, while S100a10 interacts with both tPA and plasminogen to facilitate the conversion of plasmin (Hajjar et al. 1998; Surette et al. 2011) which finally activates the matrix metalloproteinases (MMPs) (Monea et al. 2002; Davis et al. 2001). These active MMPs along with other ECM modulators promote the cleavage of key matrix proteins, including fibronectin, laminins and collagen, which helps ECM remodelling and angiogenesis during feto-placental development (Rundhaug 2005; Wang and Khalil 2018). Our placenta proteome data displayed a downregulation of plasminogen but fails to detect the expression of tPA. Therefore, we assessed the spatial distribution of tPA in BPA exposed placental sections and also explored its co-localization with Annexin A2. Interestingly, the immuno-staining reveals a significant increase in tPA levels in both decidua and labyrinth, similar to S100a10 expression. However, the co-localization of tPA and Annexin A2 decreased significantly in placental labyrinth region whereas decidual co-localization remained unchanged due to BPA exposure. These results indicate BPA triggered elevation of S100a10 actually recruits more tPA in both decidual and labyrinth fraction of placenta that drives the conversion of plasminogen to plasmin, which is evident from reduced plasminogen abundance detected in proteomics study. However, additional experiments are required for determination of the underlying causes that reduced the tPA-Annexin A2 co-localization in placental labyrinth.

This elevated plasmin, now activates the MMPs across different placental layers promoting ECM degradation, although the literature suggests a dose dependent regulation of MMP expression in response to BPA (Biswas et al. 2026b; Ye et al. 2018; Biswas et al. 2026a; Ogushi et al. 2026). Therefore, we analyzed the status of collagen deposition and the expression of matrix proteins in the placental sections, as a readout for MMP activation. The result suggests, compromised Fibronectin expression and limited collagen deposition in placental decidua due to BPA exposure. Whereas the fibronectin in placental labyrinth displays an elevated expression which may be due to reduced tPA-Annexin A2 co-localization, which is observed in previous experiments. However, the Labyrinth region did not show any signs of collagen deposition. Therefore, the results demonstrate that elevated S100a10 in placental decidua recruits more tPA in response to bisphenol-A. These may cause increased plasmin conversion, which drives the MMP activation in decidua post BPA exposure, triggering more ECM degradation. Decidual ECM remodelling is essential for feto-maternal cross talk. These indicates a S100a10-Annexin A2 axis mediated disruption of decidual ECM remodelling which drives the functional alteration of feto-maternal interface that ultimately leads to reduced feto-placental size post bisphenol-A administration.

## 5. Conclusion

Bisphenol-A has long been associated with decidual and angiogenic impairments in placenta, although the underlying mechanism required more clarity. Therefore, this current study investigates the toxicity mechanism utilizing a gestationally BPA exposed murine model to demonstrate a global placental proteome alteration which highlights the elevated regulation of S100a10 in both EVTs and placenta. Further experimentation reveals a S100a10-Annexin A2 associated upregulation of tPA in placental decidua, which results in decidual degradation of ECM components due to BPA exposure. This impaired ECM remodelling in decidua drives functional perturbation in feto-maternal axis, that ultimately culminates in decreased feto-placental size, which is evident from phenotypic assessments (**Figure 7**). However, these functional observations need to be validated in a clinical cohort to establish a broader implication of this BPA mediated toxicity mechanism during pregnancy. Despite these limitations, the study establishes a novel molecular mechanism of decidual ECM modulation via S100a10-Annexin A2 axis, as an underlying factor for BPA toxicity, which may contribute to adverse pregnancy outcome.

**Figure 7:**
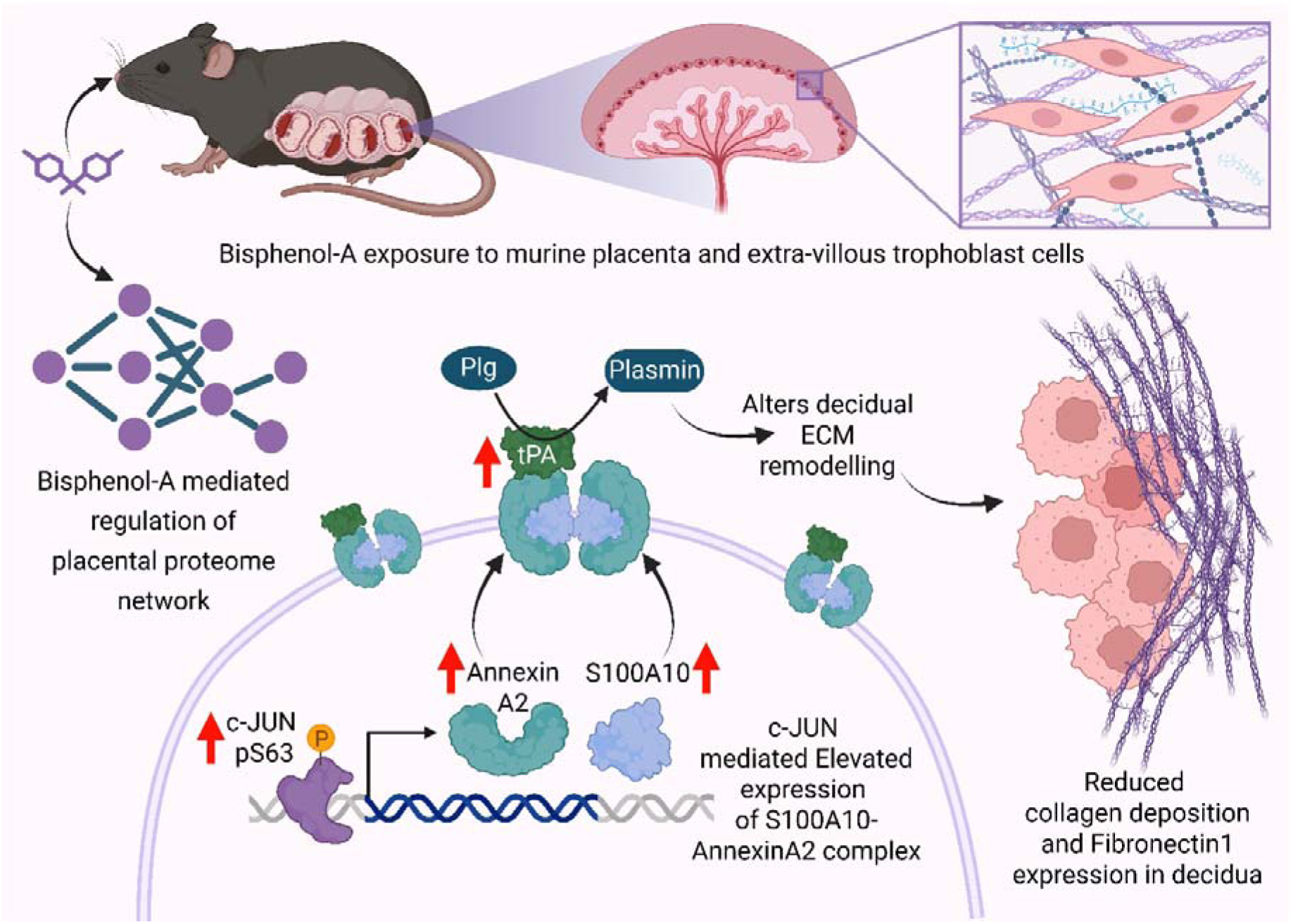
Graphical summary of the principal findings. The illustration depicts bisphenol-A mediated impairment of placenta and trophoblast proteome which highlights the perturbation of S100a10-Annexin A2 axis in placental decidua. These alterations lead to compromised ECM remodelling in placental decidua triggering functional modulations in feto-placental interface.

## Data Availability

The mass spectrometry data are available online through the ProteomeXchange Consortium via the PRIDE (https://www.ebi.ac.uk/pride/) partner repository with the data set identifier PXD077324.

## Acknowledgements

TKM acknowledges the Regional Centre for Biotechnology for its intramural research funding. We thank Dr. Pallavi khestrapal for providing us with the HTR8/SVneo cell line. We express our sincere gratitude to all the technical staff at Experimental Animal Facility at Regional Centre for Biotechnology for keeping the equipments and facilities in functional condition. We are grateful to all the members of the laboratory of functional proteomics for maintaining an excellent research environment. Somnath Mondal thanked CSIR, and Ankit Biswas thanked DBT for their fellowship.

## Author information

### Contributions

A.B. and T.K.M. conceived the project and designed the experiments. A.B. performed proteomics sample preparation, mass spectrometry data acquisition, analysis, cellular and animal experiments. S.M. performed and S.J.M. supervised the placental sectioning, immune-fluorescent staining and image acquisition. A.B. and T.K.M. wrote the manuscript. S.M. and S.J.M. edited and approved the final version of the manuscript.

### Corresponding author

Tushar Kanti Maiti, Laboratory of Functional Proteomics, Regional Centre for Biotechnology, NCR Biotech Science Cluster, Faridabad 121001, India.

## References

1. Seham A Abd El-Aleem and Lodewijk V Dekker. Assessment of the cellular localisation of the annexin a2/s100a10 complex in human placenta. Journal of molecular histology, 49(5):531–543, 2018.

2. Enoch Appiah Adu-Gyamfi, Cheryl S Rosenfeld, and Geetu Tuteja. The impact of bisphenol a on the placenta. Biology of Reproduction, 106(5):826–834, 2022.

3. Scott M Belcher, Robin B Gear, and Eric L Kendig. Bisphenol a alters autonomic tone and extracellular matrix structure and induces sex-specific effects on cardiovascular function in male and female cd-1 mice. Endocrinology, 156(3):882–895, 2015.

4. Alamelu Bharadwaj, Emma Kempster, and David Morton Waisman. The annexina2/s100a10 complex: the mutualistic symbiosis of two distinct proteins. Biomolecules, 11(12):1849, 2021.

5. Ankit Biswas, Sandhini Saha, Naman Kharbanda, and Tushar Kanti Maiti. Integrated proteomic and phosphoproteomic profiling reveals mechanisms of bisphenol a induced placental toxicity. Chemical Research in Toxicology, 2026a.

6. Ankit Biswas, Sandhini Saha, Debapriyo Sarmadhikari, Krishna Singh Bisht, Shailendra Asthana, and Tushar Kanti Maiti. Bisphenol-a mediated ubiquitinome alteration triggers ppar-alpha ubiquitination, affecting trophoblast cell migration. bioRxiv, pages 2026–05, 2026b.

7. Brad Bolon, Susan A Elmore, Wendy Halpern, and Colin G Rousseaux. Embryo, fetus, and placenta. Haschek and Rousseaux’s Handbook of Toxicologic Pathology, pages 819–917, 2025.

8. Caroline Bonnans, Jonathan Chou, and Zena Werb. Remodelling the extracellular matrix in development and disease. Nature reviews Molecular cell biology, 15(12):786–801, 2014.

9. Graham J Burton and Abigail L Fowden. The placenta: a multifaceted, transient organ. Philosophical Transactions of the Royal Society B: Biological Sciences, 370(1663):20140066, 2015.

10. David E Cantonwine, Russ Hauser, and John D Meeker. Bisphenol a and human reproductive health. Expert review of obstetrics & gynecology, 8(4):329–335, 2013.

11. David E Cantonwine, Kelly K Ferguson, Bhramar Mukherjee, Thomas F McElrath, and John D Meeker. Urinary bisphenol a levels during pregnancy and risk of preterm birth. Environmental health perspectives, 123(9):895, 2015.

12. Dominika Celar Sturm, Tadeja Režen, Nina Jančar, and Irma Virant-Klun. Bisphenol a disrupts steroidogenesis and induces apoptosis in human granulosa cells cultured in vitro. International journal of molecular sciences, 26(9):4081, 2025.

13. Revathy U Chottekalapanda, Salina Kalik, Jodi Gresack, Alyssa Ayala, Melanie Gao, Wei Wang, Sarah Meller, Ammar Aly, Anne Schaefer, and Paul Greengard. Ap-1 controls the p11-dependent antidepressant response. Molecular psychiatry, 25(7):1364–1381, 2020.

14. Wei-Chun Chou, Jyh-Larng Chen, Chung-Fen Lin, Yi-Chun Chen, Feng-Cheng Shih, and Chun-Yu Chuang. Biomonitoring of bisphenol a concentrations in maternal and umbilical cord blood in regard to birth outcomes and adipokine expression: a birth cohort study in taiwan. Environmental health, 10(1):94, 2011.

15. George E Davis, Kristine A Pintar Allen, René Salazar, and Steven A Maxwell. Matrix metalloproteinase-1 and-9 activation by plasmin regulates a novel endothelial cell-mediated mechanism of collagen gel contraction and capillary tube regression in three-dimensional collagen matrices. Journal of cell science, 114(5):917–930, 2001.

16. Reuben Blair Dodson, Paul J Rozance, Bradley S Fleenor, Carson C Petrash, Lauren G Shoemaker, Kendall S Hunter, and Virginia L Ferguson. Increased arterial stiffness and extracellular matrix reorganization in intrauterine growth-restricted fetal sheep. Pediatric research, 73(2):147–154, 2013.

17. Dana C Dolinoy, Dale Huang, and Randy L Jirtle. Maternal nutrient supplementation counteracts bisphenol a-induced dna hypomethylation in early development. Proceedings of the National Academy of Sciences, 104(32):13056–13061, 2007.

18. Eduardo Augusto Brosco Famá and Maria Aparecida Silva Pinhal. Extracellular matrix components in preeclampsia. Clinica chimica acta, 568:120132, 2025.

19. Qingyao Fu, Yuzeng Wang, Yidi Zhang, and Zhenlong Wu. Bisphenol a and bisphenol s disrupt placental epithelial-mesenchymal transition and angiogenesis via activating endoplasmic reticulum stress. Journal of Hazardous Materials Advances, 101170, 2026.

20. Cielo García-Montero, Tatiana Pekarek, Oscar Fraile-Martinez, Diego Liviu Boaru, Patricia de Castro-Martinez, Beatriz García-González, Marina Fanega-Fernández, Coral Bravo,

21. Juan A De Leon-Luis, Raul Diaz-Pedrero, et al. Late-onset preeclampsia is linked to extensive remodeling of the placental extracellular matrix. Medical Sciences, 14(3):364, 2026.

22. Katherine A Hajjar, Laura Mauri, Andrew T Jacovina, Fengming Zhong, Urooj A Mirza, Julio Cesar Padovan, and Brian T Chait. Tissue plasminogen activator binding to the annexin ii tail domain: direct modulation by homocysteine. Journal of Biological Chemistry, 273(16):9987–9993, 1998.

23. Haipeng Huang, Jiaqi Hou, Yilie Liao, Fangchao Wei, and Baoshan Xing. Polyethylene microplastics impede the innate immune response by disrupting the extracellular matrix and signaling transduction. Iscience, 26(8), 2023.

24. Sha Huang, Jiufeng Li, Shunqing Xu, Hongzhi Zhao, Yuanyuan Li, Yanqiu Zhou, Jing Fang, Jiaqiang Liao, Zongwei Cai, and Wei Xia. Bisphenol a and bisphenol s exposures during pregnancy and gestational age—a longitudinal study in china. Chemosphere, 237:124426, 2019.

25. Tina Izard. Environmental toxicants and their disruption of integrin signaling in lipid rafts. Bioessays, 47(5):e202400276, 2025.

26. Naman Kharbanda, Ankit Biswas, Arundhati Tiwari, Pragya Tailor, Sandhini Saha, Nitya Wadhwa, Ramachandran Thiruvengadam, Dinakar M Salunke, Shinjini Bhatnagar, GARBH-Ini Study Group, et al. Placental proteomics reveals an elevated level of aldo-keto reductase 1-b1, highlighting its potential role in spontaneous preterm birth. Journal of Proteome Research, 24(2):612–623, 2025.

27. Hsin-Chieh Lan, Kai-Yu Wu, I-Wen Lin, Zhi-Jie Yang, Ai-An Chang, and Meng-Chun Hu. Bisphenol a disrupts steroidogenesis and induces a sex hormone imbalance through c-jun phosphorylation in leydig cells. Chemosphere, 185:237–246, 2017.

28. Xuening Li, Qi Chen, Dan Wu, Zhe Xiao, Ce Shi, Youdan Dong, and Lihong Jia. High levels of bpa and bpf exposure during pregnancy are associated with lower birth weight in shenyang in northeast china. Chemical Research in Toxicology, 37(7):1199–1209, 2024.

29. Ning Lu, Jing Wang, Yi-hui Li, Xu-hui Fang, Yu-ting Peng, Qing-peng Luo, and Ai-hua Liao. Extracellular matrix: new insights into its role in female reproductive aging and potential therapeutic strategies. npj Aging, 2026.

30. Jiude Mao, Ashish Jain, Nancy D Denslow, Mohammad-Zaman Nouri, Sixue Chen, Tingting Wang, Ning Zhu, Jin Koh, Saurav J Sarma, Barbara W Sumner, et al. Bisphenol a and bisphenol s disruptions of the mouse placenta and potential effects on the placenta–brain axis. Proceedings of the National Academy of Sciences, 117(9):4642–4652, 2020.

31. Jenna M Mennella, Lori A Underhill, Sophia Collis, Geralyn M Lambert-Messerlian, Richard Tucker, and Beatrice E Lechner. Serum decorin, biglycan, and extracellular matrix component expression in preterm birth. Reproductive Sciences, 28(1):228–236, 2021.

32. Archana Molangiri, Saikanth Varma, Navya Sree Boga, Priti Das, Asim K Duttaroy, and Sanjay Basak. Gestational exposure to bpa alters the expression of glucose and lipid metabolic mediators in the placenta: role in programming offspring for obesity. Toxicology, 509:153957, 2024.

33. Sara Monea, Kaisa Lehti, Jorma Keski-Oja, and Paolo Mignatti. Plasmin activates pro-matrix metalloproteinase-2 with a membrane-type 1 matrix metalloproteinase-dependent mechanism. Journal of cellular physiology, 192(2):160–170, 2002.

34. Kevin Moreau, Ghita Ghislat, Warren Hochfeld, Maurizio Renna, Eszter Zavodszky, Gautam Runwal, Claudia Puri, Shirley Lee, Farah Siddiqi, Fiona M Menzies, et al. Transcriptional regulation of annexin a2 promotes starvation-induced autophagy. Nature communications, 6(1):8045, 2015.

35. Judith Elisabeth Müller, Nicole Meyer, Clarisa Guillermina Santamaria, Anne Schumacher, Enrique Hugo Luque, Maria Laura Zenclussen, Horacio Adolfo Rodriguez, and Ana Claudia Zenclussen. Bisphenol a exposure during early pregnancy impairs uterine spiral artery remodeling and provokes intrauterine growth restriction in mice. Scientific reports, 8(1):9196, 2018.

36. Saba Nikanfar, Ellen CR Leonel, Pauliina Damdimopoulou, Jodi A Flaws, and Christiani A Amorim. Effects of phthalate exposure on human ovarian extracellular matrix composition: insights from a 3d spheroid model. Environmental research, 279:121797, 2025.

37. Shoko Ogushi, Reina Abe, Takehiro Nakamura, Tsuyoshi Nakanishi, and Tomoki Kimura. Environmentally relevant low-dose bisphenol a enhances invasion of human extravillous trophoblasts with associated increases in mmp activity. Reproductive Toxicology, 109236, 2026.

38. Gillian C Okura, Alamelu G Bharadwaj, and David M Waisman. Recent advances in molecular and cellular functions of s100a10. Biomolecules, 13(10):1450, 2023.

39. Jackye Peretz, Lisa Vrooman, William A Ricke, Patricia A Hunt, Shelley Ehrlich, Russ Hauser, Vasantha Padmanabhan, Hugh S Taylor, Shanna H Swan, Catherine A VandeVoort, et al. Bisphenol a and reproductive health: update of experimental and human evidence, 2007–2013. Environmental health perspectives, 122(8):775, 2014.

40. Elise M Philips, Leonardo Trasande, Linda G Kahn, Romy Gaillard, Eric AP Steegers, and Vincent WV Jaddoe. Early pregnancy bisphenol and phthalate metabolite levels, maternal hemodynamics and gestational hypertensive disorders. Human reproduction, 34(2):365–373, 2019.

41. Muralitharan Ponniah, E Ellen Billett, and Luigi A De Girolamo. Bisphenol a increases bewo trophoblast survival in stress-induced paradigms through regulation of oxidative stress and apoptosis. Chemical Research in Toxicology, 28(9):1693–1703, 2015.

42. Divya Sangeetha Rajkumar, Gopinath Murugan, and Rajashree Padmanaban. Unraveling the interaction of bisphenol a with collagen and its effect on conformational and thermal stability. Biophysical Chemistry, 298:107026, 2023.

43. Ursula Rescher and Volker Gerke. S100a10/p11: family, friends and functions. Pflügers Archiv-European Journal of Physiology, 455(4):575–582, 2008.

44. Francesca Rossi, Stefania Luppi, Albina Fejza, Elena Giolo, Giuseppe Ricci, and Eva Andreuzzi. Extracellular matrix and pregnancy: functions and opportunities caught in the net. Reproductive biology and endocrinology, 23(1):24, 2025.

45. Joyce E Rundhaug. Matrix metalloproteinases and angiogenesis. Journal of cellular and molecular medicine, 9(2):267–285, 2005.

46. Yuriko Saiki and Akira Horii. Multiple functions of s100a10, an important cancer promoter. Pathology international, 69(11):629–636, 2019.

47. Bumpenporn Sanannam, Sasikarn Looprasertkul, Songphon Kanlayaprasit, Nakarin Kitkumthorn, Tewarit Sarachana, and Depicha Jindatip. Alteration of extracellular matrix components in the anterior pituitary gland of neonatal rats induced by a maternal bisphenol a diet during pregnancy. International Journal of Molecular Sciences, 22(23):12667, 2021.

48. Lauren Sayres, Shuhan Ji, Carla Rey Diaz, Diane Gumina, and Emily Su. The role of extracellular matrix proteins on placental angiogenesis in severe fetal growth restriction. American Journal of Obstetrics & Gynecology, 228(1):S69, 2023.

49. Amanda Nancy Sferruzzi-Perri, Jorge Lopez-Tello, and Esteban Salazar-Petres. Placental adaptations supporting fetal growth during normal and adverse gestational environments. Experimental Physiology, 108(3):371–397, 2023.

50. Yanan Sun, Menghan Sha, Yu Qin, Juan Xiao, Wei Li, Shufang Li, and Suhua Chen. Bisphenol a induces placental ferroptosis and fetal growth restriction via the yap/taz-ferritinophagy axis. Free Radical Biology and Medicine, 213:524–540, 2024.

51. Alexi P Surette, Patricia A Madureira, Kyle D Phipps, Victoria A Miller, Per Svenningsson, and David M Waisman. Regulation of fibrinolysis by s100a10 in vivo. *Blood*, The Journal of the American Society of Hematology, 118(11):3172–3181, 2011.

52. Martha Susiarjo, Isaac Sasson, Clementina Mesaros, and Marisa S Bartolomei. Bisphenol a exposure disrupts genomic imprinting in the mouse. PLoS genetics, 9(4):e1003401, 2013.

53. Sabrina Tait, Roberta Tassinari, Francesca Maranghi, and Alberto Mantovani. Bisphenol a affects placental layers morphology and angiogenesis during early pregnancy phase in mice. Journal of Applied Toxicology, 35(11):1278–1291, 2015.

54. Laura N Vandenberg, Ibrahim Chahoud, Vasantha Padmanabhan, Francisco JR Paumgartten, and Gilbert Schoenfelder. Biomonitoring studies should be used by regulatory agencies to assess human exposure levels and safety of bisphenol a. Environmental health perspectives, 118(8):1051, 2010.

55. Xi Wang and Raouf A Khalil. Matrix metalloproteinases, vascular remodeling, and vascular disease. Advances in pharmacology, 81:241–330, 2018.

56. Xiaohua Wang, Aiju Liu, Dongxia Hou, Xiaoyang Dong, Chu Chu, Weina Ju, Junhui Zhang, Yueqi Jia, Xiaoyan Yang, Yunpeng Ji, et al. Epigenetic impact of long noncoding rna lnc-adam9 on extracellular matrix pathway in preterm syndrome through down-regulation of mrna-adam9. Journal of Translational Genetics and Genomics, 7(2):126–140, 2023.

57. Zongting Wang, Ruohe An, Liang Zhang, Xiaohui Li, and Cong Zhang. Exposure to bisphenol a jeopardizes decidualization and consequently triggers preeclampsia by upregulating cyp1b1. Journal of Hazardous Materials, 486:137032, 2025.

58. Clarissa Wormsbaecher, Andrea R Hindman, Alex Avendano, Marcos Cortes-Medina, Caitlin E Jones, Andrew Bushman, Lotanna Onua, Claire E Kovalchin, Alina R Murphy, Hannah L Helber, et al. In utero estrogenic endocrine disruption alters the stroma to increase extracellular matrix density and mammary gland stiffness. Breast Cancer Research, 22(1):41, 2020.

59. Yunzhen Ye, Qiongjie Zhou, Liping Feng, Jiangnan Wu, Yu Xiong, and Xiaotian Li. Maternal serum bisphenol a levels and risk of pre-eclampsia: a nested case–control study. The European Journal of Public Health, 27(6):1102–1107, 2017.

60. Yunzhen Ye, Yao Tang, Yu Xiong, Liping Feng, and Xiaotian Li. Bisphenol a exposure alters placentation and causes preeclampsia-like features in pregnant mice involved in reprogramming of dna methylation of wnt2. The FASEB Journal, 33(2):2732, 2018.

61. Huifeng Yue, Huizhen Zhu, Xiaoyun Wu, Yuchai Tian, Jiyue Zhang, Yangcheng Hu, Xiaotong Ji, and Nan Sang. Maternal bisphenol a (bpa) exposure induces placental dysfunction and health risk in adult female offspring: insights from a mouse model. Science of the Total Environment, 958:177714, 2025.

62. Sufen Zhang, Qihan Wu, Wanhong He, Haijun Zhu, Ziliang Wang, Hong Liang, Xiaohua Ni, Wei Yuan, and Daru Lu. Bisphenol a alters jun promoter methylation, impairing steroid metabolism in placental cells and linking to sub-representative phenotypes. Gene, 941:149210, 2025.

